# Nationwide surveillance identifies yellow fever and chikungunya viruses transmitted by various species of *Aedes* mosquitoes in Nigeria

**DOI:** 10.1101/2024.01.15.575625

**Authors:** Udoka C. Nwangwu, Judith U. Oguzie, William E. Nwachukwu, Cosmas O. Onwude, Festus A. Dogunro, Mawlouth Diallo, Chukwuebuka K. Ezihe, Nneka O. Agashi, Emelda I. Eloy, Stephen O. Anokwu, Clementina C. Anioke, Linda C. Ikechukwu, Chukwuebuka M. Nwosu, Oscar N. Nwaogo, Ifeoma M. Ngwu, Rose N. Onyeanusi, Angela I. Okoronkwo, Francis U. Orizu, Monica O. Etiki, Esther N. Onuora, Sobajo Tope Adeorike, Peter C. Okeke, Okechukwu C. Chukwuekezie, Josephine C. Ochu, Sulaiman S. Ibrahim, Adetifa Ifedayo, Chikwe Ihekweazu, Christian T. Happi

**Author notes:** **Corresponding author:** (CTH).

## Abstract

**Background:** Since its reemergence in 2017, yellow fever (YF) has been active in Nigeria. The Nigeria Centre for Disease Control (NCDC) has coordinated responses to the outbreaks with the support of the World Health Organization (WHO). The National Arbovirus and Vectors Research Centre (NAVRC) handles the vector component of these responses. This study sought to identify the vectors driving YF transmission and any of the targeted arboviruses and their distribution across states.

**Methods:** Eggs, larvae and pupae as well as adult mosquitoes were collected in observational, analytical, and cross-sectional surveys conducted in sixteen YF outbreak states between 2017 and 2020. Adult mosquitoes (field-collected or reared from immature stages) were morphologically identified, and arboviruses were detected using RT-qPCR at the African Centre of Excellence for Genomics of Infectious Diseases (ACEGID).

**Results:** *Aedes* mosquitoes were collected in eleven of the sixteen states surveyed and the mosquitoes in nine states were found infected with arboviruses. A total of seven *Aedes* species were collected from different parts of the country. *Aedes aegypti* was the most dominant (51%) species, whereas *Aedes africanus* was the least (0.2%). Yellow fever virus (YFV) was discovered in 33 (∼26%) out of the 127 *Aedes* mosquito pools. In addition to YFV, the Chikungunya virus (CHIKV) was found in nine pools. Except for *Ae. africanus*, all the *Aedes* species tested positive for at least one arbovirus. YFV-positive pools were found in six (6) *Aedes* species while CHIKV-positive pools were only recorded in two *Aedes* species. Edo State had the most positive pools (16), while Nasarawa, Imo, and Anambra states had the least (1 positive pool). Breteau and house indices were higher than normal transmission thresholds in all but one state.

**Conclusion:** In Nigeria, there is a substantial risk of arbovirus transmission by *Aedes* mosquitoes, with YFV posing the largest threat at the moment. This risk is heightened by the fact that YFV and CHIKV have been detected in vectors across outbreak locations. Hence, there is an urgent need to step up arbovirus surveillance and control activities in the country.

**Author summary:** Epidemics of arboviral infections are fast becoming the norm across Africa, especially in Nigeria, where sporadic epidemics of *Aedes*-borne Yellow Fever (YF) have been on the rise since 2017. A nationwide surveillance was conducted to detect the arboviruses and the vectors transmitting them during large YF outbreaks across Nigeria. *Aedes* mosquitoes collected from 57 local government areas/outbreak locations, covering 16 out of the 36 states in Nigeria, were reared to adulthood, morphologically identified to species level and assayed using qRT-PCR to detect six epidemiologically important arboviruses (Dengue, Chikungunya (CHIKV), Zika, YF, West Nile and O’nyong nyong). Seven *Aedes* species (including *Ae. aegypti, Ae. albopictus, Ae. circumluteolus, Ae. vittatus, Ae. simpsoni* complex*, Ae. luteocephalus,* and *Ae. africanus*), were caught, with *Ae. aegypti* and *Ae. albopictus* the most widespread and predominant. Only YF and CHKV were found in mosquito pools from 9 states, with only *Ae. aegypti* pools positive for CHKV, while YF virus was found predominantly in this species and *Ae. albopictus*. This study provides a baseline for widespread distribution of the major *Aedes* vectors, and their arboviral transmission profiles, which can enhance evidence-based control measures, and strengthen nationwide surveillance systems to forestall future outbreaks of public health magnitude.

## Introduction

The encroachment of the invasive *Aedes* mosquitoes is increasing the spread of debilitating arboviral infections, such as dengue, chikungunya, and yellow fever viruses, among others (1). The invasive *Aedes* species of public health importance, have spread to all continents, except Antarctica (2) where they transmit the major arboviruses of Public Health importance. Globalization, increasing volume and pace of trade and travel, continuing urbanization and environmental challenges which include climate change are major factors driving the spread of these vectors and the diseases they transmit (3).

*Aedes aegypti* and *Ae. albopictus* are the two most medically important and invasive *Aedes* species. Together they are largely responsible for the transmission of dengue, chikungunya, yellow fever and Zika viruses around the world (4). Transmission of dengue, Chikungunya and Zika viruses in the tropics and sub-tropics is primarily by *Ae*. *aegypti*, while *Ae*. *albopictus* is mostly implicated in temperate zones and other settings (5). While dengue, chikungunya, yellow fever and Zika viruses all have sylvatic cycles involving forest mosquitoes and non-human primates, recent global outbreaks have been dominated by urban transmission by *Ae*. *aegypti* and *Ae*. *albopictus* (4). In the African settings, *Aedes aegypti* and *Ae. albopictus* are also considered the major arbovirus vectors (6). However, there are a number of minor *Aedes* mosquitoes that are epidemiologically significant secondary vectors of arboviruses. These include *Aedes africanus*, *Ae. luteocephalus*, *Ae. simpsoni* group/complex, *Ae. vittatus, Ae. metallicus*, *Ae. opok*, and *Ae. furcifer*/*taylori* group (7). Almost all of these species are found in Nigeria (8) and some have been found to habour yellow fever and dengue viruses in different parts of the country (9,10). Sylvatic dengue viruses in Africa are transmitted among non-human primates by *Ae. furcifer* and *Ae. luteocephalus*, and usually cross over to humans through biting by *Ae. furcifer* (11).

Funding gaps hugely affect the control of arboviruses in Africa. According to the (12), most countries reported a lack of adequate financial and technical support. The World Health Organization (WHO) also highlighted that thirty-four countries (72%) in the WHO African region reported that they did not have an emergency fund or a specified emergency funding mechanism for arbovirus disease outbreak response in the previous 2 years. Consequently, there is lack of training and retraining for staff involved in the surveillance and control of arboviral diseases, and even the lack of community awareness of arboviral diseases. Generally, Africa lacks systematic surveillance and reporting of many diseases, including dengue (13). Diagnostic capacity for dengue, as for virtually all causes of acute febrile illness (AFI), is limited in Africa. It is essential to understand the biology and behaviour of local vectors because these factors will influence transmission, as well as selection and design of effective control tools and strategies (14). Unfortunately, data as regards vectors and disease transmission is insufficient in most parts of Africa (6). The spread of arboviruses (chikungunya, dengue and Zika virus) in Africa is to a large extent not properly understood. Also, knowledge of the differences in the risk of transmitting these diseases and yellow fever within the sub-regions is inhibited by a lack of data (15,16).

Since the earliest recorded outbreak of yellow fever in Nigeria in 1864 (17), yellow fever has been a recurrent public health challenge in the country (18). However, there have been quiescent periods and resurgence of the disease. Yellow fever outbreaks have overlapped with outbreaks or cases of other arboviruses which are often not reported or underreported, as they have usually had a smaller impact on morbidity and mortality (19–21). More so, the similarity of signs and symptoms as well as the poor diagnosis has been a major challenge to delineating these arboviruses for proper mapping in the country. This is also a limitation to measuring the impact of the individual disease burden. There have been several reports of other arboviruses – Zika virus disease (22,23), Dengue (19,21), and Chikungunya (19,20) in several parts of Nigeria. Unfortunately, there are usually no follow-ups and measurable impacts. Sadly, these viruses may have remained in circulation for ages, unnoticed.

The resurgence of yellow fever in Nigeria started in Ifelodun Local Government Area (LGA) of Kwara State in 2017 (24). Since the year 2017 yellow fever has been reported in all 36 states and Federal Capital Territory (FCT) with 32 states reporting at least 1 confirmed case (25). Between September 2017 and December 2021, there were a total of 14,272 suspected cases from 759 (98.0%) LGAs across all the states in Nigeria. Of these 14,272, 702 cases were confirmed from the reference laboratories.

Hence, in line with the National yellow fever outbreak response strategy, we carried out a nationwide entomological surveillance to investigate the arboviruses in circulation and the *Aedes* species that transmit them across the country between the years 2017 and 2020. This will also provide information on vector distribution and composition, as well as guide the relevant authorities to formulate policies that will bring about prevention and control of the vectors and their diseases.

## Methods

### Study area

The study was carried out in fifty-seven (57) Local Government Areas (LGAs) from sixteen states in Nigeria (Fig 1). Most of the areas are rural settlements where farming is the major occupation. Nigeria is located mainly within the lowland humid tropics, characterized by high temperatures of up to 32°C in the coastal south and up to 41°C in the North (26). Nigeria is characterized by 6 ecological zones, from South to North: the Mangrove Swamp, Freshwater Swamp, Rainforest, Guinea Savanna, Sudan Savanna, and Sahel Savanna (27). The climate varies from very wet typical in coastal areas, south of Nigeria, with an annual rainfall greater than 3500mm to dry in the Sahel region in the North West and North-Eastern parts, with annual rainfall below 600mm per annum (ref), with huge climatic variations depending on the regions, with the climate becoming drier along latitudinal gradient from south to north. There are two seasons – rainy (April to October) and dry seasons (November to March) characterized by the harmattan. The harmattan season in Nigeria begins in November/December and ends in February.

**Fig. 1.**
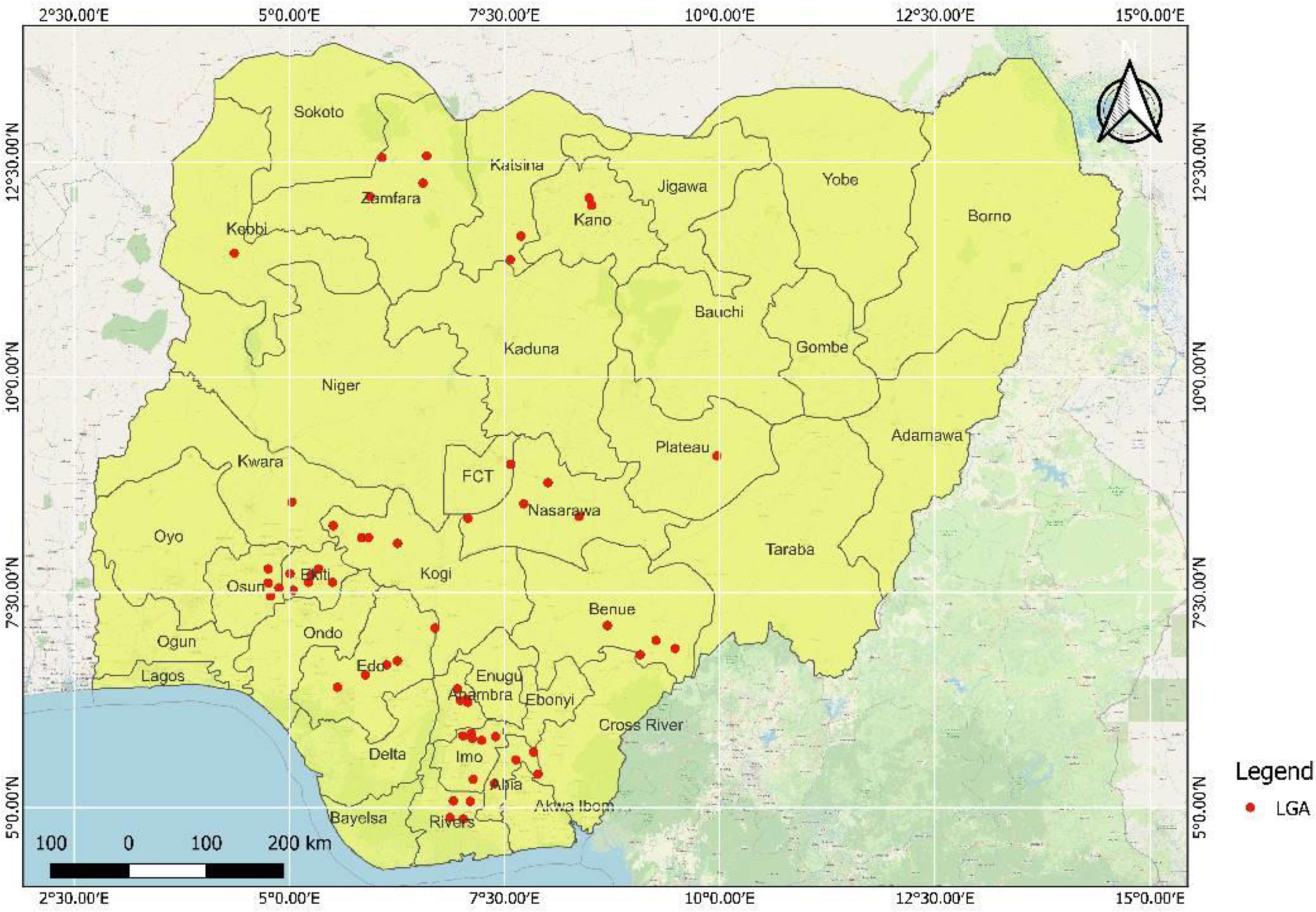
Map of Nigeria showing LGAs sampled in different states.

### Mosquito collection

Entomological approaches used in the survey included the use of ovitraps, larval survey, modified Human Landing Catch (mHLC), BG-Sentinel Trap and CDC Light-Trap. Each of the visits/responses to the YF outbreak lasted between eight and ten days.

### Use of ovitraps

Each ovitrap consists of a cylindrical cup (about 500ml in volume, containing about 200ml of water), lined with a white ribbon (cloth) on which mosquitoes laid eggs just above the water level. In each community, twenty (20) ovitraps were placed on the ground under shades, around houses and nearby bushes. The traps were collected and examined for the presence of eggs after 48 h. The ribbons were room-dried and positive ribbons (eggs on them) were separated from the negative ones. All positive and negative ribbons were pooled separately and soaked for the hatching of eggs. The negative ribbons were soaked to clear any doubt of missed egg(s).

### Larval survey

Using the house of the index case as the centre, larval sampling was carried out in houses within 300-meter radius. This activity targeted the immature stages of domestic and peri-domestic breeders among the *Aedes* species (7). Artificial containers in and around various houses were sampled for larvae and pupae of the vectors. Plant axils (natural habitats) which might harbour immature stages (larvae and/or pupae) of *Aedes* species around human dwellings were also sampled. Where possible, the entire water content of a container was emptied into a bowl and the immature stages were picked using pipettes. These were then introduced into labelled containers and reared into adults for morphological identification.

### Modified human landing catch (mHLC)

This method was carried out in accordance with (7), though modified by covering most parts of the body. Two individuals performed this across outbreak locations. Adult collections were performed in the mornings between 7 am and 10 am and then evenings between 4 pm and 8 pm in the same location, over a period of two to three days. Biting activities of most *Aedes* species peaked during these periods. Mosquitoes were caught using mouth aspirators or test tubes as soon as they landed to bite. Those collected with mouth aspirators were transferred to the test tubes. The test tubes were quickly plugged with cotton wool and transferred into cold boxes with ice packs.

### Adult collection using BG-Sentinel trap (Biogents Sentinel Trap)

This trap targets day blood-seeking female mosquitoes (7). It is used with a proprietary lure (BG-Lure®). Two BG-Sentinel traps with lures were set outdoors at each location and allowed to run for 12 hours (between 7am and 7pm). At the end of the period, the traps were collected and examined for the presence of adult mosquitoes.

### Adult collection using CDC light/UV trap

To investigate the nocturnal/crepuscular activities of some *Aedes* species, two of these traps were set outdoors in front of human dwellings between 6pm and 6am for two to three nights, in areas with outbreaks. At the end of the period, the traps were collected and examined for the presence of adult mosquitoes.

### Morphological identification and preservation of mosquitoes

Adult mosquitoes collected in the field and those reared to adults from the immature stages were chilled to death or knocked down using Ethyl Acetate. The specimens were morphologically identified on chill tables using the taxonomic keys of (28), (29) and (30). They were then introduced into well-labelled Eppendorf tubes containing RNAlater and stored at −20^0^C.

### Identification of yellow fever virus using qRT-PCR

Mosquitoes were pooled according to species, sex and location. Each pool contains 1-26 mosquitoes. Vector pools were homogenized in 1ml of cooled Dulbecco’s Modified Eagle Medium (DMEM) (composition-500ml DMEM High Glucose (4.5g/I) with L-Glutamine), 1ml Penicillin-Streptomycin, 15 ml Fetal Calf Serum (FCS) 3% and 5ml Amphotericin B) and 500ml of Zirconia beads (Firma Biospec: 2.0 mm, Cat. No 1107912). The contents were macerated using the Qiagen Tissuelyser LT for 10 min and then centrifuged at 4,500 x g for 15 min. Viral RNA was extracted from supernatant using Qiagen viral QIAamp mini kit and qRT-PCR was used to screen flaviviruses (YFV, WNV, DENV and ZIKV) and alphaviruses (CHIK and ONNV).

SYBR green RT-qPCR was performed on a Roche LightCycler 96; samples were run in triplicate and called positive after showing amplification on all replicates with amplification in the nuclease-free water used as non-template control. Briefly, 3 μL of RNA was used per reaction as a template for amplification, and this sample was added to 7 μL of reaction mixture containing 1.32 μL of H_2_O, 5 μL of Power SYBR master mix, 0.08 μL of 125X reaction mix and 0.3 μL sense and anti-sense primers.

Real-time RT-qPCR amplification was carried out for 45 cycles at 48 °C for 30 min, 95 °C for 10 min, 95 °C for 15 sec, and 60 °C for 30 sec. Temperatures for the melt curves were 95 °C for 15 sec, 55 °C for 15 sec and 95 °C for 15 sec for all previously published primers used in this study (31–35). The list of these primers is provided in Table 1 below.

**Table 1.**
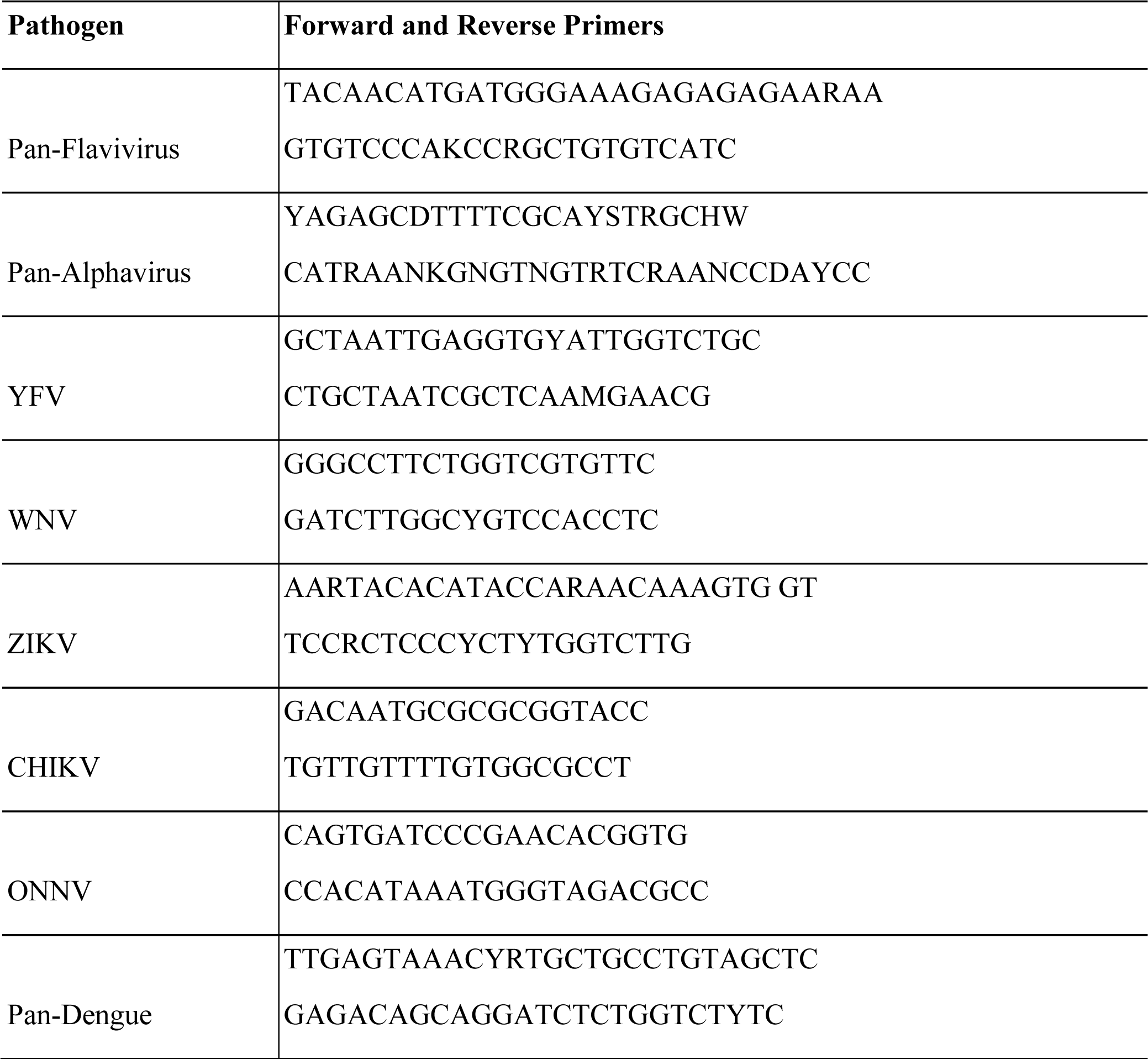
Primer sequences for the arboviruses.

### Data analyses

From larval survey, the Breteau Index (BI; number of positive containers per 100 houses), Container Index (CI; percentage of all containers with water that are larvae and/or pupae positive) and the House Index (HI; percentage of houses with at least one positive container) were estimated. All data were analyzed using SPSS version 26. Data were presented in bar chart(s), pie charts and tables. Chi-square was used to determine whether differences in number between different mosquito species are significant or not. R package was used to determine infection rates of *Aedes* mosquitoes in each state according to CDC PooledInfRate (Version 4.0) software.

## Results

### Composition, distribution, and diversity of *Aedes* species

A total of 7 *Aedes* species (*Ae. aegypti, Ae. albopictus, Ae. circumluteolus, Ae. vittatus, Ae. simpsoni* complex*, Ae. luteocephalus,* and *Ae. africanus*) were collected during the YF outbreak Rapid Response carried out in 16 states (Table 2a). However, there were nineteen (19) distorted and unidentified *Aedes* (Stegomyia) mosquitoes. *Aedes aegypti* (1,012, 42%) and *Ae. albopictus* (765, 31.8%) were clearly the two most abundant (*P* < 0.001) of all the *Aedes* species (2,406) collected. However, significantly more *Ae. aegypti* was collected than *Ae. albopictus* (*P* < 0.001). *Ae. africanus* is the least abundant of all the *Aedes* species collected (3, 0.13%).

**Table 2.** *Aedes* species composition and abundance using mHLC, emergence from larval survey and adult trap catches across states.

| State | *Ae. aegypti* | *Ae. albopictus* | *Ae. africanus* | *Ae. luteocephalus* | *Ae. simpsoni complex* | *Ae. circum-luteolus* | *Ae. vittatus* | Un-identified | Total |
| --- | --- | --- | --- | --- | --- | --- | --- | --- | --- |
| **(a) Composition and abundance of *Aedes* mosquitoes collected using mHLC across states** |  |  |  |  |  |  |  |  |  |
| Abia | 1 | 28 | 0 | 1 | 0 | 1 | 1 | 0 | 32 |
| Anambra | 12 | 44 | 0 | 0 | 1 | 0 | 0 | 0 | 57 |
| Imo | 3 | 7 | 1 | 0 | 0 | 0 | 0 | 0 | 11 |
| Benue | 60 | 7 | 0 | 4 | 0 | 0 | 0 | 0 | 71 |
| Kwara | 3 | 0 | 2 | 1 | 0 | 0 | 0 | 0 | 6 |
| Nasarawa | 1 | 0 | 0 | 0 | 0 | 0 | 0 | 0 | 1 |
| Plateau | 2 | 0 | 0 | 0 | 0 | 0 | 0 | 0 | 2 |
| Ekiti | 0 | 3 | 0 | 0 | 0 | 0 | 0 | 0 | 3 |
| Edo | 28 | 37 | 0 | 2 | 14 | 34 | 1 | 1 | 117 |
| Osun | 136 | 56 | 0 | 0 | 0 | 0 | 2 | 0 | 194 |
| Rivers | 0 | 41 | 0 | 0 | 0 | 0 | 0 | 0 | 41 |
| **(b) Composition and abundance of *Aedes* species collected through larval survey (Emergence)** |  |  |  |  |  |  |  |  |  |
| Abia | 64 | 56 | 0 | 0 | 5 | 0 | 0 | 0 | 125 |
| Anambra | 435 | 415 | 0 | 0 | 0 | 0 | 0 | 0 | 850 |
| Benue | 4 | 8 | 0 | 0 | 0 | 0 | 2 | 0 | 14 |
| Kwara | 86 | 0 | 0 | 0 | 0 | 0 | 0 | 0 | 86 |
| Nasarawa | 83 | 0 | 0 | 0 | 0 | 0 | 0 | 0 | 83 |
| Ekiti | 29 | 4 | 0 | 0 | 0 | 0 | 13 | 0 | 46 |
| Edo | 18 | 3 | 0 | 0 | 0 | 0 | 11 | 0 | 32 |
| Rivers | 1 | 13 | 0 | 0 | 0 | 0 | 0 | 0 | 14 |
| Katsina | 5 | 0 | 0 | 0 | 0 | 0 | 0 | 0 | 5 |
| Zamfara | 2 | 0 | 0 | 0 | 0 | 0 | 0 | 0 | 2 |
| **(c) Composition and abundance of *Aedes* species collected through adult trapping** |  |  |  |  |  |  |  |  |  |
| Abia | 7 | 7 | 0 | 0 | 1 | 0 | 0 | 0 | 15 |
| Anambra | 14 | 15 | 0 | 0 | 0 | 2 | 0 | 0 | 31 |
| Benue | 9 | 2 | 0 | 0 | 0 | 0 | 0 | 0 | 11 |
| Edo | 9 | 0 | 0 | 2 | 2 | 506 | 1 | 2 | 522 |
| Nasarawa | 0 | 0 | 0 | 0 | 0 | 0 | 0 | 1 | 1 |
| Rivers | 0 | 19 | 0 | 0 | 0 | 0 | 0 | 0 | 19 |

*Aedes albopictus* was by far the most abundant of all the *Aedes* species collected in the three Southeastern states of Imo, Abia, and Anambra, followed by *Ae. aegypti.* There was a statistically significant difference (*P* < 0.001) between the number of *Ae. albopictus* and the number of the rest of *Aedes* species in Abia and Anambra states and none between them (*P* > 0.05) in Imo state. In the North central states (Benue, Nasarawa, Kwara, and Plateau), *Ae. aegypti* was significantly more abundant (*P* < 0.001) than any other *Aedes* species in Benue state alone. However, no significant difference in abundance was recorded in the difference between *Aedes* species in Nasarawa, Kwara and Plateau states.

In the Southwest (SW) and South-South (SS) Zones, Edo State was the highest in species composition (all 7 *Aedes* species were collected) and abundance (*P* < 0.001). Ekiti and Osun had 3 species each, while Rivers had 2 species. *Aedes circumluteolus* was the most abundant species collected (*P* < 0.001) in the two regions (though almost exclusively from one community in Edo State), followed by *Ae. aegypti* (221), *Ae. albopictus* (176). *Ae. luteocephalus* was the least in abundance. *Aedes aegypti* was highest in abundance (*P* < 0.001) than all other *Aedes* species in Osun State, while *Ae. albopictus* was significantly higher (almost exclusively collected) in number than *Ae. aegypti* in Rivers state. *Ae. circumluteolus* was the most abundant (P < 0.001) *Aedes* species in Edo 1 (first visit to Edo State), whereas in Edo 2 (second visit to Edo State), there was no significant difference in the abundance of the different *Aedes* species collected.

*Aedes* mosquitoes that emerged from larvae or pupae that were collected during larval survey were recorded only in ten out of sixteen states namely: Abia, Anambra, Benue, Edo, Ekiti, Katsina, Kwara, Nasarawa, Rivers and Zamfara states (Table 2b). Using this method five species of *Aedes* (*Ae. aegypti, Ae. albopictus, Ae. simpsoni* complex*, Ae. vittatus* and *Ae. luteocephalus*) were collected from the ten states*. Aedes aegypti* was the highest in abundance (*P* < 0.001), followed by *Ae. albopictus (P* < 0.001). *Ae. simpsoni* complex (5) and *Ae. vittatus* (1) were the least in abundance. Three species of *Aedes* were collected in Benue (*Ae. aegypti, Ae. albopictus* and *Ae. vittatus*), Abia (*Ae. aegypti, Ae. albopictus* and *Ae. simpsoni* complex), Ekiti (*Ae. aegypti, Ae. albopictus* and *Ae. vittatus*) and Edo (*Ae. aegypti, Ae. albopictus* and *Ae. vittatus*) states. *Aedes aegypti* and *Ae. albopictus* were collected in Anambra and Rivers states, while only *Ae. aegypti* was collected from Katsina, Nasarawa, Kwara and Zamfara states. Thus, *Ae. aegypti* was collected in all the ten states in which larval sampling yielded results, while *Ae. albopictus* was recorded in six out of these ten states. There was no significant difference in the abundance of *Ae. aegypti* and *Ae. albopictus* obtained through larval survey in Anambra and Abia states (*P* = 0.493). In Rivers state, only one male *Ae. aegypti* mosquito emerged from collections, using the same method. All others were *Ae. albopictus*.

Adult trapping method was deployed and it yielded results in Abia, Anambra, Benue, Edo, Nasarawa and Rivers states (Table 2c). Vectors collected included *Ae. circumluteolus, Ae. aegypti, Ae. albopictus, Ae. simpsoni* complex*, Ae. vittatus, Ae. luteocephalus* and unidentified *Aedes* species. *Aedes circumluteolus* was the highest in abundance compared to the other *Aedes* species collected from Edo 1 (P < 0.001) and those collected from all the other states put together (P = 0.002). While only A*e. albopictus* was collected using adult trapping method in Rivers state, no significant difference was recorded in the abundance of the *Aedes* species in the five remaining states. Five *Aedes* species (*Ae. aegypti, Ae. luteocephalus, Ae. simpsoni* complex*, Ae. circumluteolus* and *Ae. vittatus*) were collected in Edo state; three species each were collected in both Abia (*Ae. aegypti, Ae. albopictus* and *Ae. simpsoni* complex) and Anambra (*Ae. aegypti, Ae. albopictus* and *Aedes circumluteolus*) states; two species (*Ae. aegypti* and *Ae. albopictus*) were collected in Benue state and one species (*Ae. albopictus*) each was collected in both Nasarawa and Rivers states.

### Epidemic risk indices and breeding preferences across states

Results of the epidemic risk indices (Fig 2) show that House and Breteau indices were higher than the standard threshold across the states, except in Kogi. Container index was also higher than standard risk threshold in all localities except Kogi and Zamfara states.

**Fig 2.**
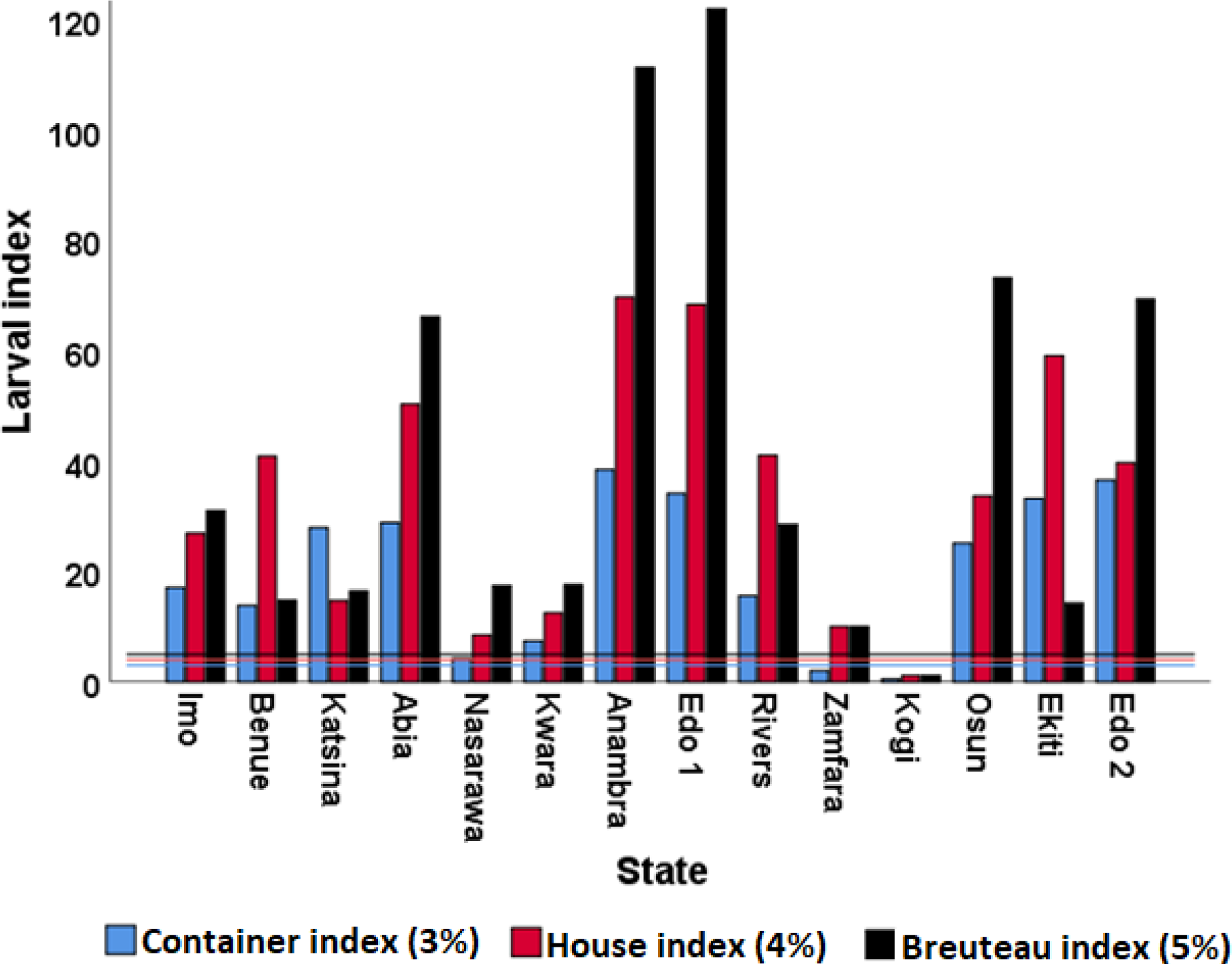
Epidemic risk indices across states.

Results of container typology revealed that the predominant breeding site of *Aedes* mosquitoes in the study area is plastic containers, followed by tyres, earthen wares, metal containers and the least in abundance is rock pools (Fig 3). This pattern was observed across the states from the northern to the southern parts of the country. However, mosquitoes collected from Nasarawa and Osun states showed striking preference for tyres.

**Fig. 3.**
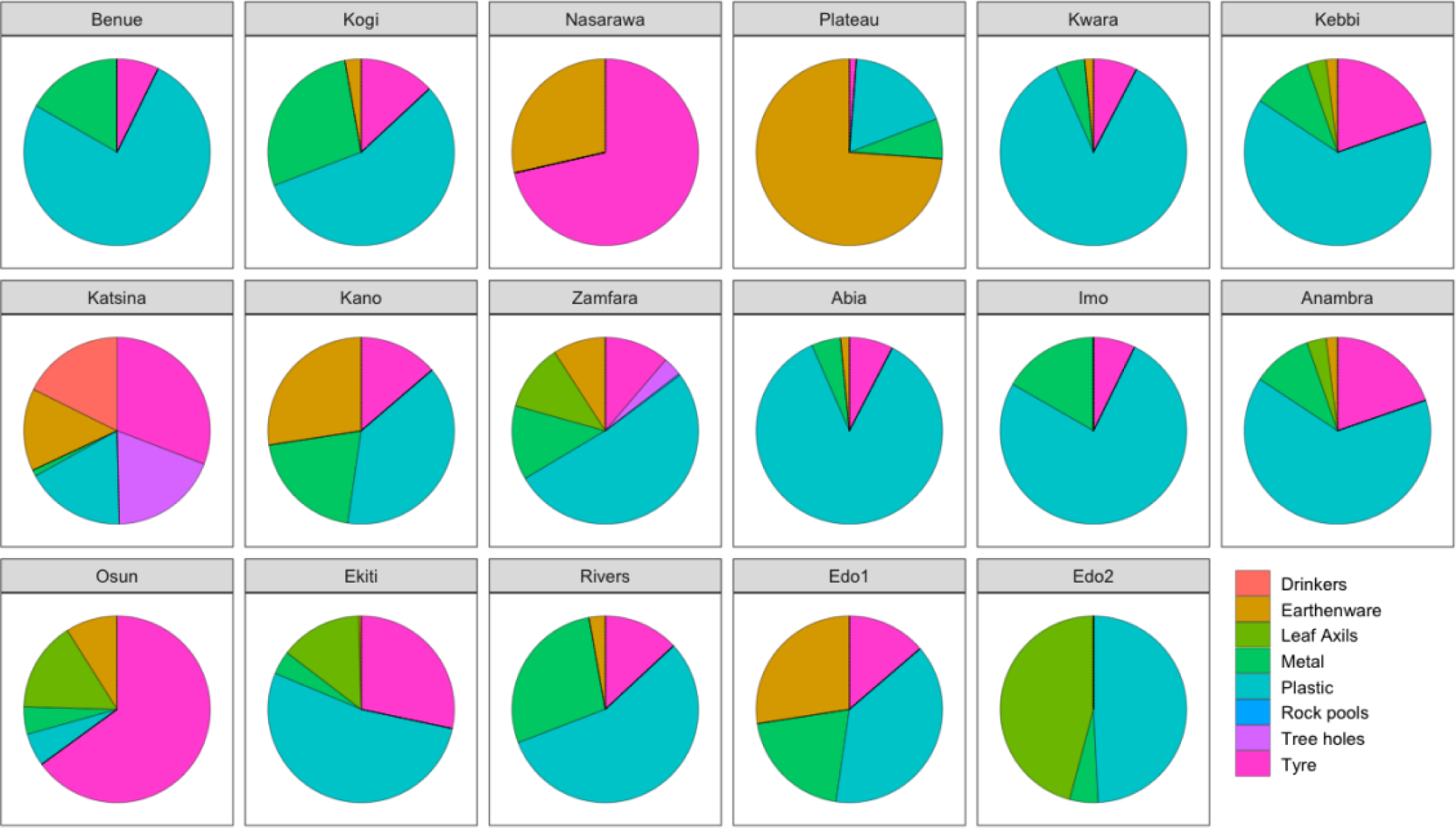
Container typology for breeding of *Aedes* mosquitoes.

### Mosquito infectivity with arboviruses across the states

Of the six arboviruses screened, only yellow fever virus (YFV) and chikungunya virus (CHIKV) were detected (Tables 3 and 4). Nine (82%) out of eleven states had *Aedes* mosquito pools positive for the YFV. Two states (18%) only, had no YFV-positive mosquitoes – Plateau and Ekiti states. However, in Edo state, roughly 64% (14/22) of mosquito pools was positive for the YFV.

**Table 3.** *Aedes* species YFV and CHIKV infection rates across southern states.

| State | Species | Number Collected | YFV |  |  |  |  | CHIKV |  |  |  |  |
| --- | --- | --- | --- | --- | --- | --- | --- | --- | --- | --- | --- | --- |
|  |  |  | Pools tested | Positive Pool | Infection rate (P) | 95% CI Lower | 95% CI Upper | Pools tested | Positive Pool | Infection rate (P) | 95% CI Lower | 95% CI Upper |
| Abia | *Ae. aegypti* | 72 | 6 | 1 | 2.320 | 0.1387 | 11.78 | 6 | 0 | 0.000 | 0.000 | 6.85 |
|  | *Ae. albopictus* | 91 | 7 | 2 | 3.405 | 0.6296 | 11.74 | 7 | 0 | 0.000 | 0.000 | 4.80 |
|  | *Ae. simpsoni* | 6 | 2 | 1 | 85.167 | 5.5058 | 434.85 | 2 | 0 | 0.000 | 0.000 | 163.59 |
|  | *Ae. spp* | 15 | 6 | 0 | 0.000 | 0.000 | 32.45 | 6 | 0 | 0.000 | 0.000 | 32.45 |
| **Sub total** |  | **184** | **21** | **4** | **0.190** |  |  | **21** | **0** | **0** | **0** | **0** |
| Anambra | *Ae. aegypti* | 461 | 2 | 0 | 0.000 | 0.000 | 2.32 | 2 | 0 | 0.000 | 0.000 | 2.32 |
|  | *Ae. albopictus* | 474 | 2 | 1 | 1.078 | 0.0699 | 7.20 | 2 | 0 | 0.000 | 0.000 | 2.26 |
|  | *Ae. simpsoni* | 1 | 1 | 0 | 0.000 | 0.000 | 793.45 | 1 | 0 | 0.000 | 0.000 | 793.45 |
|  | *Ae. circumluteolus* | 2 | 1 | 0 | 0.000 | 0.000 | 545.52 | 1 | 0 | 0.000 | 0.000 | 545.52 |
| **Sub total** |  | **938** | **6** | **1** |  |  |  | **6** | **0** | **0** | **0** | **0** |
| Imo | *Ae. aegypti* | 3 | 1 | 1 | 1 | NA | NA | 1 | 0 | 0.000 | 0.000 | 404.88 |
|  | *Ae. albopictus* | 7 | 2 | 0 | 0.000 | 0.000 | 141.97 | 2 | 0 | 0.000 | 0.000 | 141.97 |
|  | *Ae. africanus* | 1 | 1 | 0 | 0.000 | 0.000 | 793.45 | 1 | 0 | 0.000 | 0.000 | 793.45 |
| **Sub total** |  | **11** | **4** | **1** |  |  |  | **4** | **0** |  |  |  |
| Edo | *Ae. aegypti* | 55 | 4 | 3 | 19.849 | 5.5223 | 73.27 | 4 | 2 | 10.659 | 2.0254 | 40.06 |
|  | *Ae. albopictus* | 40 | 7 | 4 | 18.920 | 6.3491 | 49.79 | 7 | 0 | 0.000 | 0.000 | 10.88 |
|  | *Ae. simpsoni* | 16 | 5 | 3 | 51.671 | 14.3538 | 154.11 | 5 | 0 | 0.000 | 0.000 | 37.29 |
|  | *Ae. circumluteolus* | 540 | 2 | 1 | 0.946 | 0.0613 | 6.32 | 2 | 0 | 0.000 | 0.000 | 1.98 |
|  | *Ae. luteocephalus* | 4 | 1 | 0 | 0.000 | 0.000 | 325.85 | 1 | 0 | 0.000 | 0.000 | 325.85 |
|  | *Ae. spp* | 3 | 1 | 1 | NA | NA | NA | 1 | 0 | 0.000 | 0.000 | 408.88 |
|  | *Ae. vittatus* | 13 | 2 | 2 | NA | NA | NA | 2 | 0 | 0.000 | 0.000 | 79.14 |
| **Subtotal** |  | **670** | **22** | **14** | 0.636 |  |  |  |  |  |  |  |
| Ekiti | *Ae. aegypti* | 29 | 12 | 0 | 0.000 | 0.000 | 9.53 | 12 | 0 | 0.000 | 0.000 | 9.53 |
|  | *Ae. albopictus* | 7 | 3 | 0 | 0.000 | 0.000 | 111.10 | 3 | 0 | 0.000 | 0.000 | 111.10 |
|  | *Ae. vittatus* | 13 | 2 | 0 | 0 | 0 | 79.14 | 2 | 0 | 0.000 | 0.000 | 79.14 |
| **Subtotal** |  | **49** | **17** | **0** |  |  |  | **17** | **0** | **0** | **0** | **0** |
| Rivers | *Ae. aegypti* | 1 | 1 | 0 | 0.000 | 0.000 | 793.45 | 1 | 0 | 0.000 | 0.000 | 793.45 |
|  | *Ae. albopictus* | 73 | 5 | 2 | 6.182 | 1.1598 | 22.15 | 5 | 0 | 0.000 | 0.000 | 7.78 |
| **Subtotal** |  | **74** | **6** | **2** |  |  |  | **6** | **0** | **0** | **0** | **0** |
| Osun | *Ae. aegypti* | 136 | 12 | 1 | 0.613 | 0.0359 | 3.03 | 12 | 1 | 0.613 | 0.0359 | 3.03 |
|  | *Ae. albopictus* | 56 | 1 | 1 | NA | NA | NA | 1 | 0 | 0.000 | 0.000 | 27.77 |
|  | *Ae. vittatus* | 2 | 1 | 1 | NA | NA | NA | 1 | 0 | 0.000 | 0.000 | 545.52 |
| **Subtotal** |  | **192** | **14** | **3** | 0.2142 |  |  | **14** | **1** |  |  |  |
| **Total** |  | **2118** | **90** | **25** |  |  |  |  |  |  |  |  |

**Table 4.**
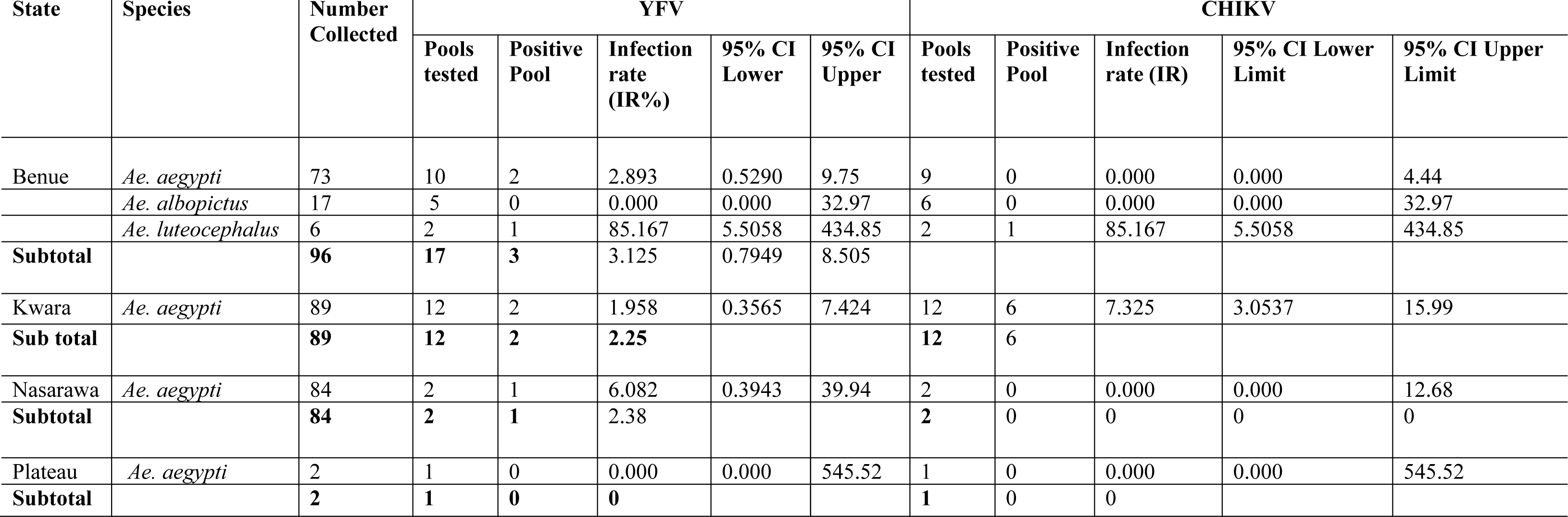
*Aedes* species YFV and CHIKV infection rates across northern states.

Of the 7 *Aedes* mosquito species collected across the country, only *Ae. africanus* recorded no YFV-positive pool (Table 3). Mosquito pools’ infectivity with YFV was highest in *Ae. aegypti* (11), followed by *Ae. albopictus* (10), *Ae. simpsoni* complex (4), *Ae. vittatus* (3) and the rest had one pool each, positive for the YFV. *Uranotaenia* species had 100% infection (1/1) with YFV. Also, YFV-positive pools were found in unidentified species of *Aedes* mosquitoes. The infection rates of the vectors by states and species only, are also shown in Supplementary information 1 and 2, respectively.

Only four (Kwara, Edo, Osun, and Benue states) out of eleven states had mosquito pools positive for the Chikungunya virus. Kwara had 50% (6/12) of its pools positive, Edo had 9.1% (2/22), Osun had 7.1% (1/14) and Benue had 6.0% (1/17) of its pools positive for the CHIKV.

Only two *Aedes* species (*Ae. aegypti* and *Ae. luteocephalus*) had CHIKV-positive pools. This represents 29 % of the total number of *Aedes* species collected. *Ae. aegypti* had 16.1% (9/56) of its pools positive for CHIKV while *Ae. luteocephalus* had 33.3% (1/3) of its pools positive for the virus. However, a logistic regression analysis showed no relationship between infectivity of mosquitoes with CHIKV and species of mosquitoes (OR = 1). This is shown in Supplementary information 3.

## Discussion

This study investigated the presence of different arbovirus vectors (mosquitoes) and their roles in the transmission of arboviruses during yellow fever outbreaks in Nigeria between 2017 and 2020. Flaviviruses (Yellow fever, West Nile, dengue and Zika viruses) and alphaviruses (Chikungunya and O’nyong nyong viruses) were screened across sites and mosquito pools.

Yellow fever and several other arboviruses are endemic to Africa, including Nigeria. These viruses are maintained and transmitted within and between animal and human populations by several *Aedes* species. Findings from this study revealed the presence of several established vectors of yellow fever and other arboviruses across the different ecological and geopolitical zones of Nigeria.

Our finding revealed a decrease of the *Aedes* species richness from the south to the north but also some disparity according to individual state data recorded during previous studies in the same zone. The rainforest Southeast (Abia, Anambra and Imo states) as well the South-South (Edo and Rivers states) Geopolitical Zones seem to be the richest ecozone in terms of *Aedes* mosquito species diversity, accounting for all 7 species (*Aedes aegypti*, *Ae. albopictus*, *Ae. africanus*, *Ae. luteocephalus*, *Ae. simpsoni* complex, *Ae. circumluteolus*, and *Ae. vittatus*) collected as adult mosquitoes in the course of the study. The seven species were collected in Edo State, which was found to have the greatest diversity of all the states. On the other hand, all the adults collected from mHLC in Rivers State were *Aedes albopictus*. Apart from one male *Aedes aegypti* emergence from larval sampling, *Aedes albopictus* was almost exclusively the only species collected across all sites visited in Rivers State. This is contrary to the findings of (36), who found only *Aedes aegypti* from larval surveys in their study area.

It is noteworthy that, *Ae. circumluteolus* was dominant in Agenebode community of Edo State. The team discontinued collection during mHLC, because of the unbearable number of *Aedes circumluteolus* alighting to bite. Hence, only the CDC UV Light trap was deployed thereafter.

The unusual dominance of the biting population of *Ae. circumluteolus* in Agenebode is to our knowledge, the first record of such an occurrence in Africa, as not so much information of this species is recorded, due to its relatively usual small numbers in the course of surveillance. (8), in a study that spanned seven years across several states in Nigeria, only recorded 40 *Aedes circumluteolus* from 7 of the 15 states where surveillance was carried out. On the contrary, (37) recorded large numbers of *Aedes circumluteolus* in Tongaland from vegetation. However, they reported very small numbers biting humans.

The two South-West states (Osun and Ekiti) sampled in the course of this study reported three species (*Aedes aegypti*, *Ae. albopictus* and *Ae. vittatus*). These species were also collected at their immature stages from larval sampling within and around human dwellings. This data agrees with the findings of (38), in Oshogbo metropolis (Osun State), who also found the three *Aedes* species. However, this contrasts the work of (8), who collected five *Aedes* species (including *Aedes aegypti* and *Ae. albopictus*) in a study carried out in the neighbouring Ondo state in 2010. In addition to the three vectors, we recorded in the South-West, they also collected *Ae. luteocephalus* in their 2011 study in Ogun State.

Of the five *Aedes* species (*Aedes aegypti*, *Ae. albopictus*, *Ae. luteocephalus*, *Ae. vittatus* and *Ae. africanus*) collected across the *Guinea Savannah* – *North Central (NC) Geopolitical Zone,* 3 species was present in each of Kwara, Nasarawa, and Plateau states, while Benue State recorded four species. This is in line with the findings of (8) who reported eight and six different *Aedes* species from Benue State in studies carried out in years 2007 and 2010, respectively. Also, other studies have found up to five or more *Aedes* species in Benue State (10,39).

Considering the very dry nature of the Sahel Savannah – North West Geopolitical Zone and the timing of the responses, collections were only made in Katsina and Zamfara states, where just *Ae. aegypti* was collected. This agrees with the findings of (8) which reported *Ae. aegypti* almost exclusively in these northern states. A recent study (40) also corroborated this finding, as they found only *Ae. aegypti* in Kano State.

Larval sampling in the course of this study was majorly carried out in domestic and peri-domestic natural and artificial containers. Findings from this study revealed that a majority of containers infested, were from artificial domestic and peri-domestic containers. (41) and (42) also had similar reports in their studies in Garoua, Cameroon and Enugu, Nigeria, respectively. In addition, our study also found used/discarded tyres with immature stages of mosquitos across several states. This agrees with the findings of (43) and (44) who reported large proportions of their collections from used/discarded tyres.

From the larval sampling emergence across the states, *Aedes aegypti* and *Ae. albopictus* preferred household water storage containers (though they were occasionally collected from tyres and plant axils), while *Ae. vittatus* and *Ae. simpsoni* complex preferred tyres and plant axils, respectively. This partly agrees with the findings of (42), who exclusively collected *Aedes aegypti* from water storage containers, despite collecting *Ae. albopictus* as adults from the same locations in Enugu and (38) who reported exclusive preference of *Ae. albopictus* for tyres, in a study in Oshogbo. The predilection of *Ae. vittatus* for used/discarded tyres was also reported by (45) in Ethiopia. However, a report on the preference for used/discarded tyres by *Ae. aegypti* in the same study, contrasts with our findings. Surprisingly, *Ae. luteocephalus*, an established tree-hole breeder, emerged from domestic and peri-domestic container collections in Benue State. Though a rare occurrence, (46) also found *Ae. luteocephalus* and *Ae. africanus* breeding in household water storage containers in Udi Hills of Enugu.

Findings from this study revealed that all the larval indices were higher than the standard threshold in all but one (Kogi) of the states where the indices were calculated. The high larval indices observed across the study sites agree with findings within (24) and outside (43) Nigeria.

*Aedes aegypti* is generally referred to as urban yellow fever/arbovirus vector. This is largely due to its adaptation to and the overwhelming dominance of urban areas. However, the only two urban areas where surveillance was carried out in our study suggest otherwise. *Aedes albopictus* was the only biting adult *Aedes* species collected in various parts of Port Harcourt (Rivers State – South-South Zone). Also, in the Southeast Zone, it was by far the dominant adult collection in the Awka area (Anambra State). This finding suggests a gradual replacement of the once-dominant *Aedes aegypti* by *Ae. albopictus*, in some of Nigeria’s urban areas. The work of (8), showed a steady increase in the density of *Aedes albopictus* in an urban area - Enugu, Nigeria, (between years 2008 and 2014) and an eventual dominance between the years 2012 and 2014, corroborates the findings of this work. The dominance of *Ae. albopictus* in Enugu State seems to be spreading beyond the city into suburbs and rural areas now. In contrast, (40), in their study in Northern Nigeria (a hotter and dryer Zone with a much fewer diversity of *Aedes* species), found only *Ae. aegypti* from collections made from cities in Kano and Bauchi states, Nigeria. However, this may be largely in part due to the high temperatures and low relative humidity (typical of northern Nigeria) which (47) reported to result in high mortality of *Ae. albopictus* eggs. This vector is also reported to establish in areas with annual mean temperatures of between 5°C and 28.5°C (2,48) and a relative humidity of 52% and above (49). Findings from several countries in Africa and Asia have reported dominance of *Ae. aegypti* within the cities, as *Ae. albopictus* dominates the suburban/rural areas (44,50). However, (50) reported an incident in which the invasive *Ae. albopictus* was gradually replacing the native *Ae. aegypti* in the Republic of the Congo, while (51) and (52) reported a seeming replacement of *Ae. aegypti* by *Ae. albopictus* in urban and peri-urban areas of Yaoundé, Cameroon. These reports also corroborate our findings. Hence, it is suggestive that with a strong surveillance system, more such findings could be revealed in various parts of Africa.

The vaccination coverage (54%) of yellow fever in Nigeria is not sufficient to provide herd immunity and prevent outbreaks (53). This is more reason why there have been repeated outbreaks across the country, as there are infected mosquitoes in various states amidst a large naïve population. YFV was detected in mosquito pools collected from nine of eleven states. Overall, over thirty-five percent of the pools were positive for the virus. Interestingly, the virus was detected in mosquito pools from four of the six Geopolitical Zones of the country. This is also a reflection of the different ecozones of Nigeria. The other two Zones (drier northern part of the country) not represented had no mosquito representations, as the areas were too dry by the time surveillance/response was carried out.

*Aedes aegypti* and *Ae. albopictus* were the most abundant across the study sites/states. These species seem to be the two most likely vectors responsible for the ongoing transmission cycles of yellow fever in Nigeria, accounting for about 64% of all positive pools. It is worthy of note that though *Ae. aegypti* had the most positive pools overall, only about 19% of its pools assayed for viruses were positive. On the other hand, over 31% of the *Ae. albopictus* pools assayed were positive for YFV. This is particularly interesting as yellow fever-infected wild populations of *Ae. albopictus*, had never been reported (54) until the year 2018, when (55) reported the first-ever yellow fever-infected population of *Ae. albopictus* in wild settings (http://www.iec.gov.br/portal/descoberta/). To our knowledge, this is only the second time wild populations of this vector are found infected with YFV, globally. This finding is particularly important as this invasive species, *Ae. albopictus* (56) has been shown to be a potent transmitter of yellow fever in laboratory settings (54). Generally, our finding on other vectors infected with YFV agrees with the Jos, Plateau 1969 yellow fever outbreak, where *Ae. luteocephalus* was found to be chiefly involved in the transmission of the virus (9). This is also similar to more recent findings in Benue State, where *Ae. aegypti* (39) and then *Ae. aegypti* and *Ae. luteocephalus* were found to be positive for the yellow fever virus (57). It is noteworthy that *Ae. luteocephalus* may have been involved in the transmission of yellow fever in the North Central region of Nigeria for over forty years.

*Ae. aegypti* which evidently has a wider distribution in Nigeria (8) had positive pools in seven of the nine states (both southern and northern states) where positive pools were recorded, while *Ae. albopictus* recorded positive pools in five states (southern states only). *Aedes albopictus* seems to be much more established in the southern parts (wetter and more forested) of Nigeria. (40) in a recent study in Kano and Bauchi states (North West and North East, Nigeria), didn’t find any *Ae. albopictus*. Such a recent finding buttresses earlier reports indicating that, *Ae. albopictus* is not so well established in the northern parts of the country. As a result, it is not surprising that no *Ae. albopictus* pool was positive for the virus in the three North Central states from where pools were assayed for the virus.

Unlike yellow fever, there are no approved vaccines or drugs for Chikungunya virus (CHIKV) disease. Being a more tropical disease with high morbidity and low mortality, it is globally under-diagnosed and under-reported. Our study reports the presence of CHIKV in mosquitoes for the first time, from yellow fever outbreak locations in Nigeria. This is suggestive of the possibility of co-infection of the diseases in those locations. CHIKV was detected in mosquito pools collected from four of eleven states from where pools were made. Overall, about eight percent of the pools were positive for the virus. The virus was detected in mosquito pools from three of the six Geopolitical Zones of Nigeria. While there were no mosquito pools from the North East Zone of the country, (19) had reported the circulation of CHIKV in febrile patients visiting hospitals in Borno State. This is an indication that with proper surveillance and diagnosis, the virus may be detected across Nigeria.

Chikungunya Virus was detected in *Aedes aegypti* and *Ae. luteocephalus* pools from four states. The virus was detected in *Aedes aegypti* pools from three (Edo, Kwara and Osun states) of the four states and *Ae. luteocephalus* in only one state (Benue). Both species are frequently found infected by CHIKV in nature in Senegal (58,59). This finding agrees with that of (20), which confirmed the circulation of CHIKV among febrile patients in Kwara State, through serological studies. Also, given the distribution of *Ae. albopictus* in Nigeria, (52) suggested the involvement of the vector in the circulation of the virus in the country. This contrasts our finding, as CHIKV was not detected in any of the *Ae. albopictus* pools in this work. However, it does not rule out the possibility of the involvement of this vector, as our work did not cover the entire country. More so, there is evidence that CHIKV circulation in Africa for the last two decades is restricted to the East-Central-South-African (ECSA) genotype, with increased fitness to the *Aedes* species circulating the CHIKV: the E1:A226 V variant associated with increased fitness to *Ae. albopictus* and the mutations E1:K211E and E2:V264A are associated with increased fitness to *Ae. aegypti* (52). Hence, the massive presence of these two vectors in Nigeria and the fact that CHIKV was detected in *Ae. aegypti*, is a call for urgent action against possible outbreak of the disease in Nigeria.

## Conclusion

The persistent outbreaks of yellow fever in Nigeria, as well as the detection of both YFV and CHIKV in *Aedes* mosquitoes from this study, underscore the endemicity of these arboviruses in the country. Given the low vaccination coverage and the distribution and diversity of established vectors across the country, the risk of transmission of these arboviruses (including the vaccine-preventable YFV) remains very high. As a result, there is an urgent need to strengthen the surveillance system of arboviruses and their vectors across Nigeria for effective control.

## Acknowledgements

We are grateful to all community stakeholders whose support ensured we were able to seamlessly carry out this work.

## Funding statement

This work received support from ACEGID laboratory and TED’s Audacious Project, including the ELMA Foundation, MacKenzie Scott, and the Skoll Foundation. This work was supported by grants from the National Institute of Allergy and Infectious Diseases (https://www.niaid.nih.gov), award numbers U01HG007480 to C.T.H), NIH-H3Africa (https://h3africa.org) award number U54HG007480. The World Bank grants projects ACE-019 and ACE-IMPACT. This work was also supported by the Rockefeller Foundation (Grant #2021 HTH), the Africa CDC through the African Society of Laboratory Medicine [ASLM] (Grant #INV018978), and the Science for Africa Foundation. Field surveillance was supported by the World Health Organization. The funders had no role in study design, data collection and analysis, decision to publish, or preparation of the manuscript.

## Supporting Information

S1 Table. YFV and CHIKV infection rates across states

S2 Table. YFV and CHIKV infection rates by *Aedes* species

S3 Table. Infectivity of *Aedes* mosquitoes with YFV and CHIKV

S4 Table. Logistic regression analysis for YFV and CHIKV infections of mosquitoes across all the states

S5 Table. Breeding sites of the mosquitoes

S6 Table. Mosquito infection with arboviruses

S7 Table. Infection rate of the mosquitoes

## Notes

### Competing Interest Statement

The authors have declared no competing interest.

